# Midbrain modulation of conditioned learning

**DOI:** 10.1101/2020.02.07.936963

**Authors:** Giorgio Rizzi, Zhuoliang Li, Norbert Hogrefe, Kelly R. Tan

## Abstract

The firing activity of striatal cholinergic interneurons (CINs) is a neural correlate of conditioned learning. However, its circuitry is not entirely described. Here we report that midbrain GABA and Glutamate neurons discriminate auditory cues and encode the association of a predictive stimulus with a footshock. Through their mono- and di-synaptic inputs via the thalamic-parafasicular sub-nucleus onto CINs, they contribute to processing fear learning.

## Introduction

In the Basal Ganglia, the peculiar firing activity of dorso-medial CINs is largely recognized as a key component in fear learning processing^1–3^. In terms of input circuitry involved in such neuronal encoding we propose that midbrain GABA and Glutamate cells may play a significant role. Indeed, GABA neurons have been shown to mono-synaptically inhibit accumbal cholinergic interneurons and optogenetic activation of this pathway led to enhanced performance in a fear conditioning task^4^. Furthermore, both GABA and Glutamate cells are excited by aversive stimuli^5–8^.

## Results

To validate our hypothesis, we first imaged the activity of midbrain cells while mice performed a classical fear conditioning task (Figure 1, Movie 1). AAV1-CamK2a-GCaMP6m was injected in the midbrain and a GRIN-lens implanted above (Figure 1A-1C). Eight to ten weeks post-surgery, mice underwent a fear conditioning paradigm (Figure 1D). They exhibited low baseline contextual freezing levels (Figure 1E) and froze significantly more to CS+ as compared to CS−; hence learning that CS+ is predictive of a foot-shock (FS, Figure 1F, 1G). As a proxy for cellular activation, calcium transients were concomitantly imaged. Cells responding to the cues were sparse and distinct (Figure 1H). All imaged neurons were profiled as responsive if their activity to a stimulus was statistically different from baseline. Throughout the learning phase, the proportion of excited neurons to CS+ increased as did the CS+ evoked-freezing levels (Figure 1I-1N, 1G). In addition, the amplitude of the calcium responses to CS+ was significantly larger from those of CS− (Figure 1O) and across days CS+ responses were discriminated from baseline (Figure 1P). On average, midbrain neurons discriminated CS+ from CS− and consolidated this since the proportion of cells engaged in positive discrimination increased over learning (Figure 1Q). We then examined the FS responses; a large majority of imaged cells were excited and their responses also discriminated from baseline (Figure S1A-S1E). Finally, we assessed the ability of midbrain cells to associate the predictive cue with the FS and analyzed the similarity between CS+ and FS responses; applying the Euclidean distance analysis. Within a conditioning session the distance significantly decreased across pairings (Figure S1F-S1H), indicating that the value of CS+ and FS became similar. Most interestingly, it also decreased throughout the learning (Figure S1I-S1J) suggesting that these neurons associate CS+ with FS such that CS+ becomes a predictive stimulus. Although the calcium indicator was expressed in all cell-type, we assumed that the imaged cells were GABA or Glutamate neurons because they exhibited excitation to FS (dopamine neurons show inhibition^5,7,9^). In addition, counterstaining of the midbrain tissue reveals that cells located within the imaging field of view are tyrosine hydroxylase (TH) negative, so not dopaminergic (Figure S1K).

**Figure 1:**
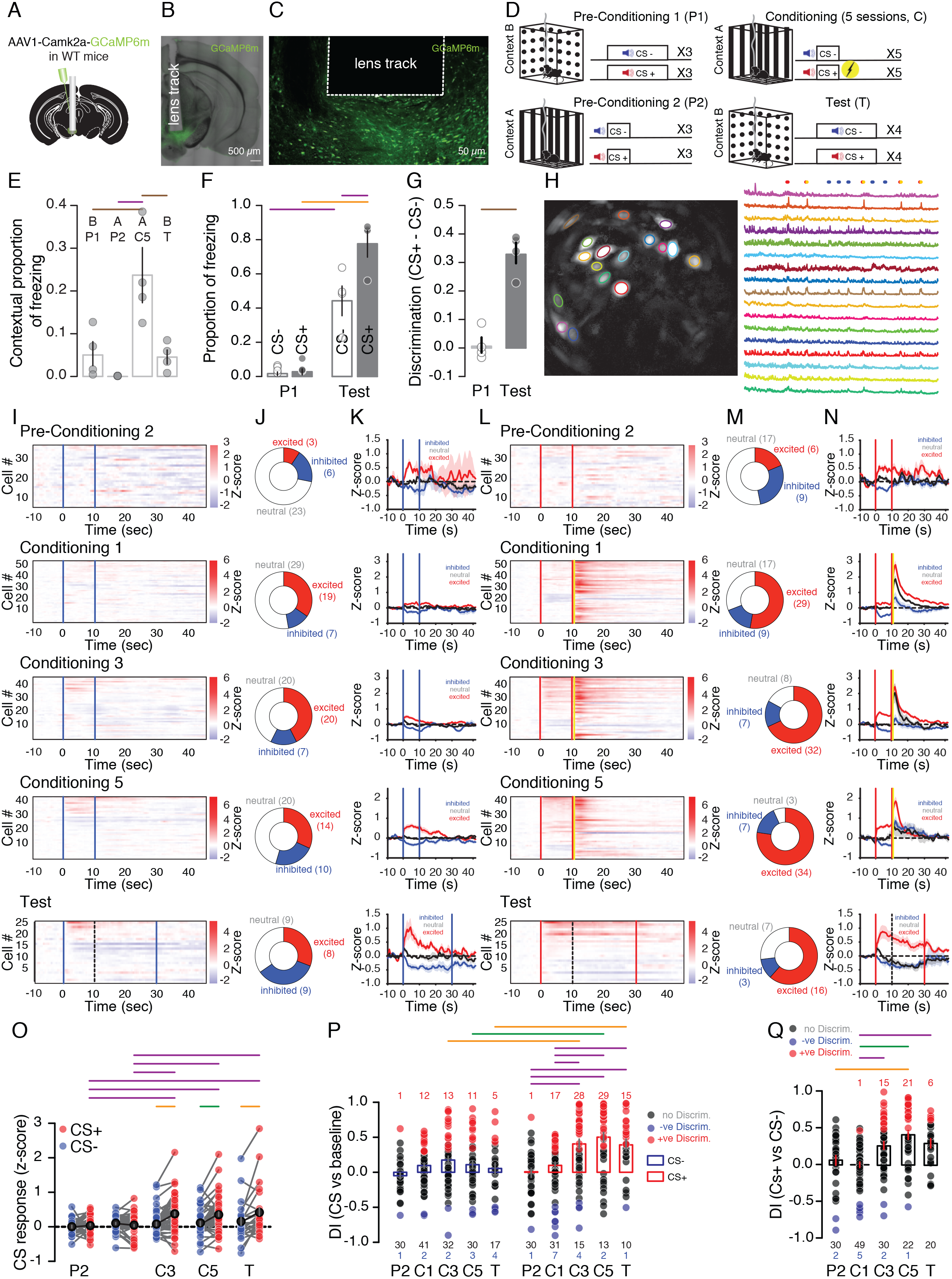
Midbrain neurons associate CS+/shock. **(A)** Surgery. (**B)** Confocal image of the midbrain. **(C)** High magnification from B. **(D)** Fear-conditioning paradigm. **(E)** Contextual freezing. **(F)** Tone freezing. **(G)** Tone discrimination. **(H)** Field of view maximal-projection image and calcium transients during C5. **(I)** Z-score heatmaps of average calcium transients to CS− (blue bars, dashed line depicts the end of the analysis time window. **(J)** Quantification. **(K)** Z-score of the average calcium transients to CS−. **(L)** Z-score heatmaps of calcium transients to CS+ and foot-shock (red and yellow bars). **(M)** Quantification. **(N)** Z-score curves of calcium transients to CS+. **(O)** Z-score of the tone responses. **(P)** Discrimination between tones and baseline. **(Q)** Discrimination between tones. *, **, ***, ****.

Because similarly to CINs, midbrain cells have the ability to predict the FS; to further confirm their role in CIN’s function; we next verified that they form functional synapses between each other (Figure 2). A transsynaptic rabies tracing approach (injection of AAV2/1-DIO-Ef1α-TVA-T2A-CVS11G and EnvA-coated SADΔG rabies into the dorso-medial striatum -DMS-in ChAT-cre mice) was first used to comprehensively explore CINs’ afferent diversity (Figure 2A). Mouse brains were either processed for tissue clearing (Figure 2B, Movie S2) or sectioned and immuno-stained (Figure 2C, 2D). Strong rabies-EGFP labelling was observed in cortical and thalamic regions^3,10^. Notably, a cluster of neurons was identified in the midbrain (Figure 2B-2D, Movie S2). These rabies labelled cells were counter-stained against TH and 96% were TH-negative (Figure 2D), corroborating our previous assumption that our neuronal population of interest is composed of GABA and glutamate cells. This is further substantiated by our fluorescent *in-situ* hybridization experiments showing that these neurons do not co-express Vgat/TH (1.4%) or Vglut2/TH (1.8%) (Figure S2).

**Figure 2:**
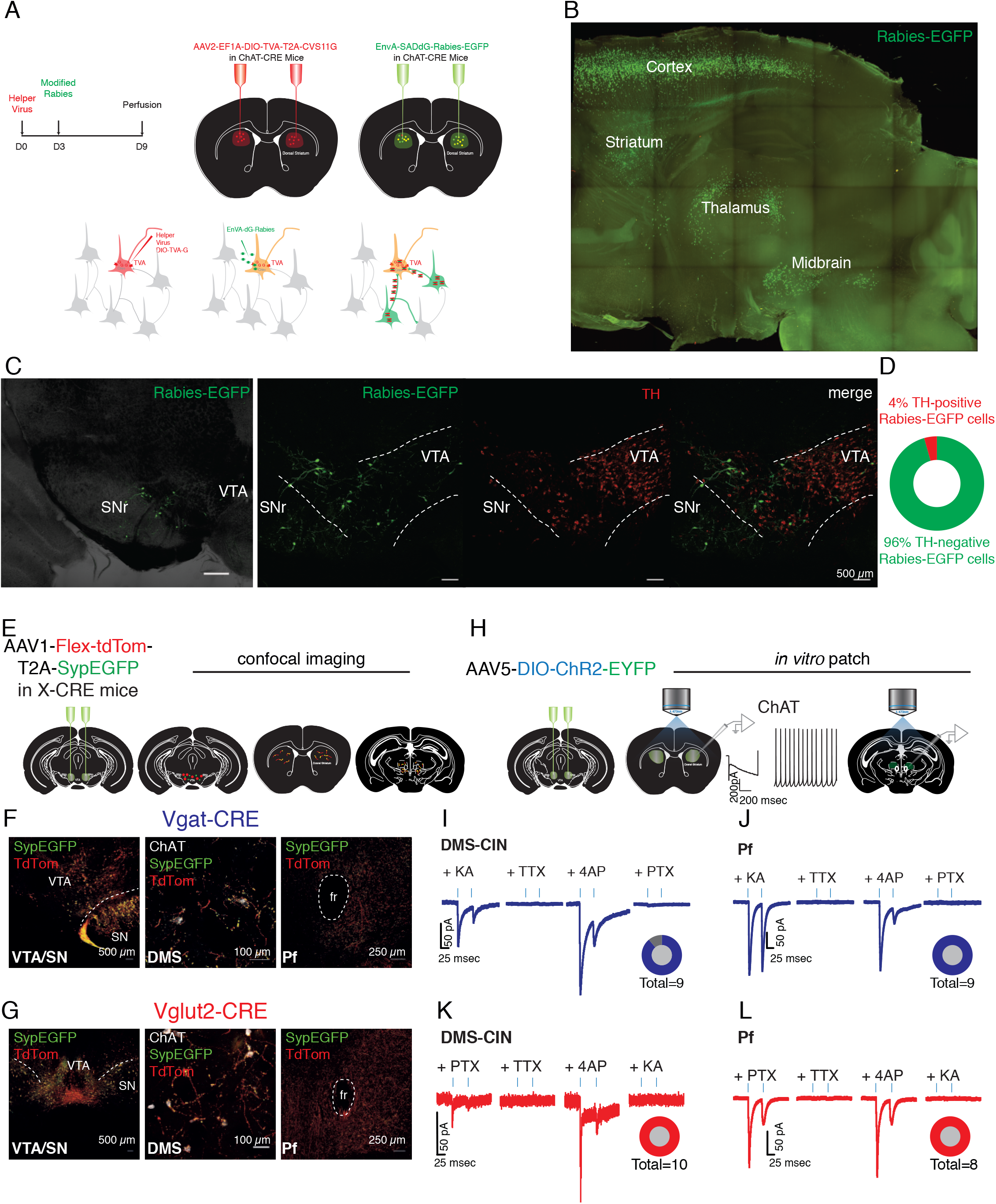
Midbrain GABA and Glutamate neurons mono-synaptically contact CINs and Pf cells. **(A)** Experimental design. **(B)** Maximal projection light-sheet image of a cleared hemisphere showing CIN inputs. **(C)** Confocal images of a midbrain containing-slice locating midbrain inputs and counter-stained against TH. **(D)** Quantitative pie chart. **(E)** Mapping experimental design. **(F)** and **(G)** Confocal images from Vgat-CRE and Vglut2-CRE mouse, respectively of Td-tomato and synaptophysin-EYFP expression in the midbrain, DMS and Pf. **(H)** Functional experimental design. **(I)** Monosynaptic inhibitory currents from midbrain-Vgat-positive cells. **(J)** Pf neurons are also modulated by GABAergic monosynaptic inputs. **(K)** Light-evoked monosynaptic excitatory currents in CINs from Vglut2-CRE mice preparations. **(L)** Monosynaptic glutamatergic currents are also recorded in Pf cells.

We also assessed the CIN to midbrain pathway via anterograde tracing and injected AAV1-phSyn1-Flex-tdTomato-T2A-Syp-EGFP in the midbrain of Vgat- and Vglut2-CRE mice (Figure 2E). In both cases, dense labelling was observed in the DMS; with synaptophysin signal apposing ChAT-immuno-positive soma. Interestingly the Pf, previously reported as necessary in CIN’s function^11^ was also heavily innervated (Figure 2F, 2G). Finally, we explored the functionality of each sub pathway, expressing ChR2-eYFP in midbrain Vgat- and Vglut2-positive cells and recording *in vitro* light evoked currents from CINs and Pf cells (Figure 2H). In all cases, robust monosynaptic currents were recorded as challenged by the successive bath-applications of TTX and 4AP (Figure 2I-2L). CINs were identified based on their intrinsic electrophysiological properties (capacitance: 58.7 ± 3.3 pF, Vm: −49.3 ± 1.4 mV, spontaneous firing and Ih current upon hyperpolarization of Vm, Figure 1F, Bennet 2000). In parallel, we verified and confirmed the monosynaptic Pf to DMS pathway applying the same mapping and functional approach (Figure S3).

So far, our data report that GABA and Glutamate midbrain neurons encode CS-FS associations and form mono- and di-synaptic contacts (via the Pf) onto CINs.

Our final step was to provide experimental evidence that these sub-pathways are instrumental in fear conditioned learning. We opted for a loss of function optogenetic manipulation of GABA and Glutamate release in the DMS and the Pf. Arch-EYFP was expressed bilaterally in the midbrain of Vgat- and Vglut2-CRE mice and optic fibers implanted in the DMS or the Pf (Figure 3A, 3B). We first verified *in-vitro* that 532nm-green light triggered currents and robust hyperpolarization, as well as blockade of induced-firing in infected neurons (Figure 3C). Mice underwent the fear conditioning task (Figure 3D). Each experimental group (Figure 3E, 3I, 3M, 3Q) showed similar low contextual freezing levels during the first conditioning and test sessions (Figure 3F, 3J, 3N, 3R) and all control mice froze more on the conditioning day 5. This was concomitant to a higher proportion of freezing to CS+ as compared to CS− on the test day (Figure 3G, 3K, 3O, 3S) and reflects association of the CS+ with the FS (Movie S3, S4).

**Figure 3:**
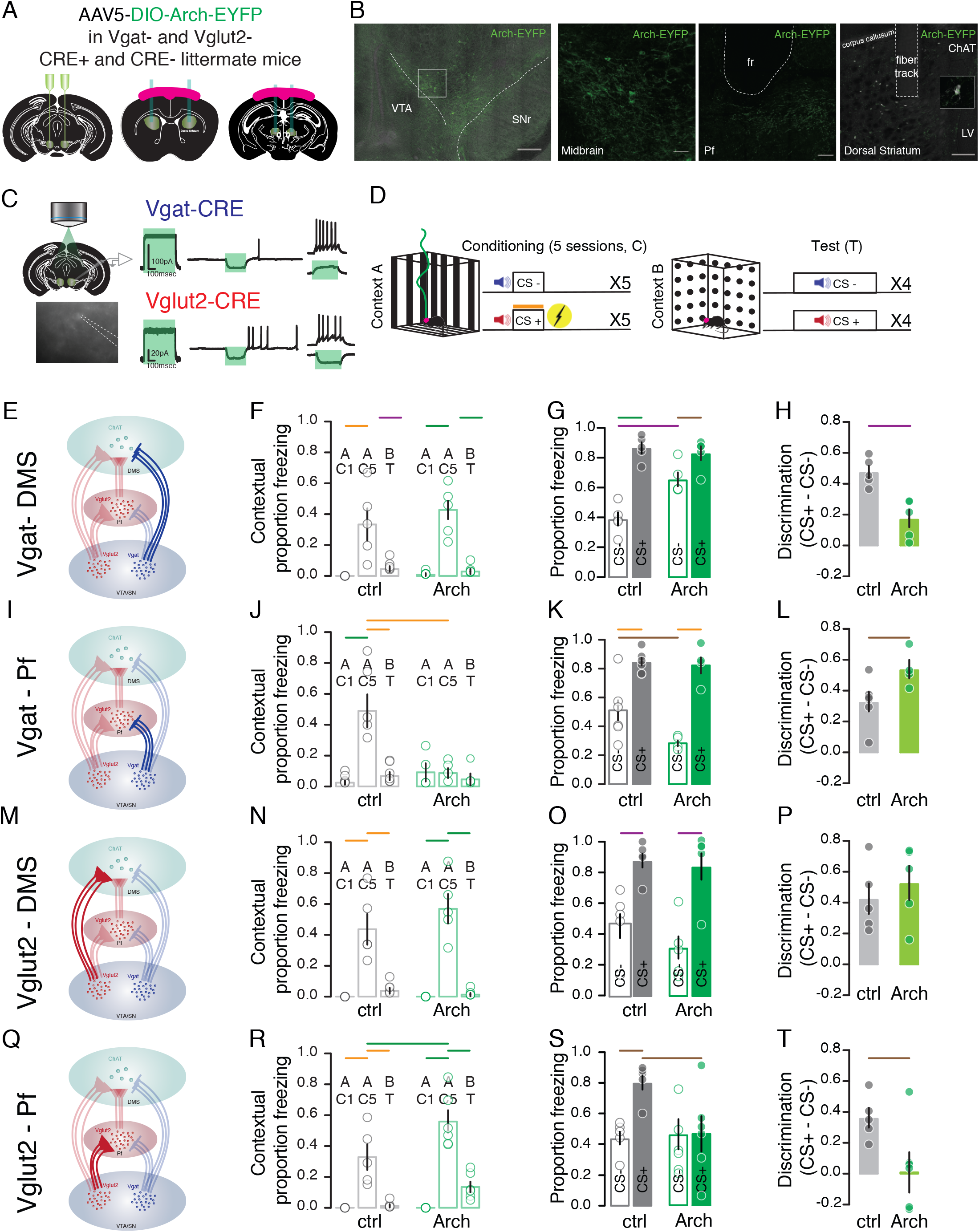
Midbrain GABA and Glutamate inputs differentially impact discrimination performance. **(A)** Surgeries. **(B)** Confocal images of the midbrain, Pf and DMS. **(C)** Optogenetic *in-vitro* validation. **(D)** Fear-conditioning paradigm. **(E)** Vgat-DMS manipulation. **(F)** Contextual freezing, **(G)** Tone freezing, **(H)** Discrimination to CSs in E. **(I)** Vgat-Pf manipulation. **(J)** Contextual freezing, **(K)** Tone freezing, and **(L)** Tone discrimination in I. **(M)** Vglut2-DMS manipulation. **(N)** Contextual freezing. **(O)** Tone freezing, **(P)** Tone discrimination, in M. **(Q)** Vglut2-Pf manipulation. **(R)** Contextual freezing, **(S)** Tone freezing to tones, **(T)** Tone discrimination in Q. *, **, ***, ****.

When GABA release was inhibited in the DMS, Arch-mice froze more to CS− as compared to controls, leading to a lower discrimination index between tones (Figure 3G, 3H). This suggests that removal of direct inhibition in the DMS from the midbrain decreases mouse discrimination abilities. When the release of GABA was inhibited in the Pf, Arch-mice exhibited less freezing to CS− hence discriminated better the two stimuli than control mice (Figure 3K, 3L). When modulating excitatory neurotransmission, inhibition of glutamate release in the DMS produced no difference between mouse groups (Figure 3O, 3P). Interestingly when the release of glutamate was inhibited in the Pf, freezing to both CS+ and CS− was low (Figure 3S). Arch-mice did not associate the CS+ with the FS (Figure 3T). These experiments confirm that each midbrain pathway, direct or indirect via the Pf, is differentially implicated in associative learning and supply CINs with specific information about behaviorally relevant events and strategies.

## Discussion

The midbrain encodes reward prediction error and learning (Schultz 1997); here we shed light onto its inhibitory and excitatory components as instrumental for CINs-mediated associative learning. Monitoring midbrain neuronal calcium activity showed that they discriminate salient stimuli and pair predictive cues with aversive outcomes, remarkably reflecting not only the behavioral freezing phenotype but also the cellular activity pattern of CINs. Although the cellular identification per se has not been assessed, the recorded neuronal activity patterns and post-hoc staining data are experimental evidence for large inclusion of GABA and Glutamate neurons rather than dopamine cells. We believe that such mirrored activity of midbrain cells and CINs is not surprising as they are wired together. Indeed, both GABA and Glutamate midbrain cell types send monosynaptic inputs onto CINs. In addition they do also contact the Pf; which itself excites CINs^11^.

Finally, our optogenetic manipulations during fear conditioning provides an undoubtable additional evidence for midbrain role in CINs aversive stimuli encoding. Inhibition of GABAergic projections to the DMS and the Pf triggered opposing behavioral performance, which based on our circuit mapping data was expected. In contrast, glutamatergic neurons showed heterogeneity; with the Pf but not DMS-projecting ones participating in the task. Manipulation of GABA- and Glutamatergic projections in the Pf induced opposing behavioral responses also in line with the circuitry we report. In-depth investigations of identified synaptic partners (simultaneously midbrain GABA/Glutamate/Pf neurons with CINs and GABA/Glutamate neurons with Pf cells) would be necessary to better understand the precise engagement of each sub-pathway in conditioned learning. Unfortunately, this requires challenging methodologies limited at the moment.

Overall, we provide additional knowledge on the circuitry underlying conditioned fear learning. Inhibitory and excitatory midbrain inputs supply CINs with information to create conditioned associations. The modulation of each pathway during learning alters different aspects of the behavioural response as a whole, suggesting they each contribute to the generation of appropriate behavioural strategies in response to external stimuli. These data may reveal valuable information, specifically when considering circuit-based therapeutical strategies in Basal Ganglia-related cognitive symptoms.

## Methods

### Animals

Male and female ChAT-CRE (B6;129S6-Chat<tm2(cre)Lowl>/J, JAX), Vglut2-Cre (B6.Cg-Slc17a6<tm2(cre)Lowl>Unc/J, UNC), Vgat-Cre (B6.Cg-Slc32a1<tm2(cre)Lowl>/J, JAX), and wildtype (WT) C57BL/6 mice were bred and genotyped in-house. All experimental procedures were approved by the Institutional Animal Care Office of the University of Basel and the Cantonal Veterinary Office under the License Number 2742.

### Surgery

#### General

Mice were anesthetized with Isoflurane (5% for induction and 1.5% for maintenance) in O_2_ (Provet/Primal Healthcare, EZ Anesthesia Systems) and placed onto the stereotaxic frame (World Precision Instruments). Lidocaine (0.2mg/g) (Steuli Pharma) was injected subcutaneously above the skull. The skin was disinfected with 70% ethanol and Betadine (Mundipharma) before drilling a hole above the target area and specific constructs were injected into the target areas using a piston operated injector (Narishige). Optic fiber and gradient-index lens implantation were carried out after injection during the same surgical session to minimize the stress to the animals. After surgical interventions, in cases of anatomical mapping or *in vitro* patch-clamp recording investigations, mice were stitched with tissue absorptive silk sutures (SABANA) and provided with temporary silicone (Smooth-On). When a GRIN-lens was implanted, a fixed headcap was built from layers consisting of super-glue (Cyberbond) and UV-light curable glue (Loctite), then secured to the skin with Vetbond tissue adhesive glue (3M). Finally, when optic fibers were implanted, super-glue and dental cement (Lang) were used to secure them to the skull via small screws. Buprenorphine (Reckitt Benckiser) pain killers were administered (0.1g/g) post-surgery as needed.

#### Optogenetic manipulations

Young adult male and female (6-8 weeks) Vgat- and Vglut2- positive mice and their CRE-negative littermates were operated on as described above, to express a CRE-dependent opsin construct (AAV5-EF1α-DIO-eArch3.0-EYFP or AAV5-EF1α-DIO-ChR2-H134R-EYFP, UNC Vector Core) bilaterally (150nl/side) in the midbrain (−3.1mm from the Bregma, + 0.5mm to the midline, and 4.25mm below the surface of the brain) and to place optic fibers (>70% light transmittance, 200m diameter 0.39NA optic fibers fixed by epoxy to 1.25mm wide 6.4mm long ceramic ferrules, Thorlabs, Sparta et al. 2010) bilaterally above either the parafasicular thalamic nucleus (Pf, −2.3mm from the Bregma, +1.15mm to the midline, and 3mm below the surface of the brain, 5 degrees away from the vertical axis) or the dorsal striatum (DMS, +0.65mm from the Bregma, + 1.7mm to the midline, and 2.5mm).

#### Calcium Imaging

Adult male and female (8-10 weeks) WT mice were surgically operated on as described above, to express a virus containing calcium indicators (AAV1.Camk2a.GCaMP6m.WPRE.SV40, Ready to Image Inscopix virus) unilaterally (200nl) in between the VTA and the SN (15 degree angle, −3.1mm from the Bregma, + 0.8mm to the midline, and 4.25mm below the surface of the brain) and to place a 600-μm-diameter (7.3mm long) gradient refractive index (GRIN) lens (Inscopix) at the same coordinates, guided by a sterile needle. The lens was then secured with UV-light curable glue (Henkel). A custom-made head bar (2cm long, 0.4cm wide, 0.1cm tall) was placed and together fixed with the GRIN lens to the brain with a combination of super-glue (Cyberbond) and dental cement (Lang). Lens clearing was assessed 4-8 weeks post infection. Mice were restrained by the implanted head bars to a fixed plexiglass running wheel (15cm diameter) and a baseplate attached miniaturized microscope (miniscope) suited for the nVoke2 system (Inscopix) was lowered to the GRIN lens until the best focal plane has been reached. Once the conditions were visually determined to be satisfactory, the mice were placed under Isoflurane anesthesia on a stereotaxic frame and the baseplate was secured to the headcap which holds the GRIN lens using a UV-light curable flowable composite glue (Kerr).

### Anatomical mapping

#### Rabies tracing

To map the monosynaptic inputs onto DMS-CINs neurons, the avian receptor restricted replication depleted modified rabies system was used (Leinweber et al., 2017). Adult ChAT-CRE mice were first injected bilaterally (300nl/side) with CRE dependent helper virus for the monosynaptic retrograde rabies system, AAV2/1-EF1α-DIO-TVA920-T2A (FMI) into the DMS, followed by EnvA-SADG-GCaMP6s into the same site three days later. The animals were sacrificed after ten days. Mice were injected intraperitoneally (ip) with pentobarbital (0.3mg/kg) (Esconarkon, Steuli Pharama AG) diluted in sterile saline and transcardially perfused with 1X phosphate buffered saline buffer (PBS) and 4% paraformaldehyde (PFA) (Sigma Aldrich) solution prepared in 1X PBS. Brains were extracted and post-fixed in 4% PFA in 1X PBS overnight then immersed in 30% sucrose in 1X PBS for at least 2 days. Sections were prepared (60 m) using a cryostat (Leica 1950 CM) and mounted for confocal imaging.

#### Anterograde tracing

To map the projections of midbrain GABAergic, glutamatergic, and dopaminergic neurons to the DMS-CINs neurons and to the Pf, Vgat- and Vglut2-CRE were injected with the CRE-dependent fluorescent construct (AAV2/1-phSyn1-FLEX-tdTomato-T2A-SypEGFP, Salk Institute) between the VTA and the SN. The CRE expressing infected cells and their processes are labeled with tdTomato and their presynaptic terminals with GFP. The animals were sacrificed after 4 to 6 weeks. Brain sections were prepared as mentioned above.

#### Retrobeads tracing

To validate the projection of the Pf to the DMS, red fluorescent microspheres (RetroBeads, Lumafluor) were injected bilaterally (150nl/side) into the DMS of adult WT mice. They were sacrificed after 10 days of expression and brain sections prepared as described above.

### *In-vitro* electrophysiology

Acute coronal slices (200μm) containing the midbrain, the DMS of the Pf were prepared using a vibrotome (Leica VT1200S) in iced-cold cutting solution (in mM: NMDG 92, KCl 2.5, NaHPO4 1.25, NaHCO_3_ 30, HEPES 20, Glucose 25, Thiourea 2, Na-Ascorbate 5, Na-Pyruvate 3, MgSO4.7H2O 10, CaCl_2_.4H2O 0.5, and N-acetyl Cystein 10, pH 7.3, Osmolarity 290-300 mOsm). Slices were incubated in ACSF solution (in mM: NaCl 92, KCl 2.5, NaH2PO4 1.25, NaHCO3 30, HEPES 20, Glucose 25, Thiourea 2, Na-ascorbate 5, Na-Pyruvate 3, MgSO4.7H_2_O 2, CaCl_2_.4H20 2, pH 7.3, Osmolarity 290-300 mOsm) at 31°C. Slices were then transferred to the recording chamber, super-fused with Ringer’s solution (in mM: NaCl 119, KCl 2.5, NaH2PO4 1.25, NaHCO3 24, Glucose 12.5, CaCl2.4H2O 2 and MgSO4.7H2O, pH 7.3 Osmolarity 290-300 mOsm) at 2ml/min bubbled with 95% O_2_ and 5% CO_2_. Neurons were visualized with an infrared (IR) camera on an Olympus scope U-TV1X-2 and whole cell patch clamp recordings (multiclamp 700B amplifier) were performed. The internal solution contained for voltage clamp recordings (in mM excitatory transmission: CsCl 130, NaCl 4, Creatine Phosphate 5, MgCl_2_ 2, Na_2_ATP 2, Na_3_GTP 0.6, EGTA 1.1 and HEPES 5, Spermine 0.1, pH 7.3, osmolarity 290 and for inhibitory transmission: KGluconate 30, KCl 100, Creatin Phosphate 10, MgCl2 4, Na2ATP 3.4, Na3GTP 0.1, EGTA 1.1 and HEPES 5, pH7.3, osmolarity 289) and for current clamp recordings (in mM: KGluconate 130, Creatin Phosphate 10, MgCl2 4, Na2ATP 3.4, Na3GTP 0.1, EGTA 1.1 and HEPES 5, pH7.3, osmolarity 289).

ChR2 stimulation was done with 2 pulses of 5msec 473nm blue light, 50msec apart with a Cool-LED pE-100 mounted on the microscope. Picrotoxin (PTX) was used at 100μM, kynureic acid (KA) at 2mM, Tetanus Toxin (TTX) at 500nM and 4-amino pyridine (AP) at 300μm.

### Histology

#### Immunohistochemistry

Midbrain or DMS-containing slices were washed (3×3 mins) with 1X Tris-buffered saline (TBS) with 0.1% Tween-20 (Sigma Aldrich), permeabilized (20mins) with 1X TBS with 0.1% Tween-20 and Triton-X100 (Sigma Aldrich), blocked (120 mins) with 1% Bovine Serum Albumin (Sigma Aldrich) in 1X TBS with 0.1% Tween-20, stained overnight with the primary antibody (Rabbit anti-TH, Sigma Aldrich T8700, 1:500 or Goat anti-ChAT, EMD Millipore AB144P, 1:200), washed, stained (120 minutes) with the secondary antibody (Goat anti-Rabbit conjugated to AlexaFlour555, Invitrogen A21429, 1:500 or Donkey anti-Goat conjugated to AlexaFlour647, abcam ab150131, 1:500), washed, mounted on glass slides (Superfrost Plus, Thermo Scientific), immersed by DAPI containing ProLong Gold Antifade mounting solution (Invitrogen) and covered with borosilicate cover glass (VWR).

#### Fluorescent in-situ hybridization (FISH)

Adult male and female wildtype C57BL/6 mice were anesthetized with pentobarbital (0.3mg/kg) (Esconarkon, Steuli Pharama AG). Brain were extracted and quickly frozen on aluminum foil over dry ice. Brains were kept overnight at −80°C. Thin 20μm fresh frozen brain sections containing the midbrain were manually prepared with the aid of a cryostat and placed onto glass slides. Selected slices were fixed for 30 minutes using 4% PFA in 1X PBS, dehydrated with increasing concentrations of ethanol (50%, 70%, 100%, 100%), surrounded by hydrophobic barriers (ImmEdge, Vector Laboratories), before proceeding with the RNAscope Fluorescent Multiplex (ACD, Biotechne) Assay to label the mRNAs for tyrosine hydroxylase (TH, ref 317621), vesicular glutamate transporter-2 (Vglut2, Slc17a6, ref319171), and vesicular GABA transporter (Vgat, Slc32a1, ref 319191). Processed slides were immersed in ProLong Diamond Antifade mounting solution (Invitrogen) and covered with borosilicate cover glass.

#### Brain clearing

To visualize the monosynaptic inputs onto DMS-CINs in 3-dimensions, a modified CUBIC-L/RA (clear, unobstructed brain imaging cocktails and computational analysis) brain clearing method (Tainaka et al., 2018) was utilized. Anatomically labeled mice were similarly transcardially perfused as described above. Immediately after perfusion, brains were washed in shaking cold (4°C) 1X PBS (3×3 mins) before being post-fixed in 4% PFA in 1X PBS under refrigeration (4°C) for 48 hours. Fixed brains were split sagittally along the midline and individually submerged into an eppendorf tube (5ml) filled with pre-heated (to 37 °C) CUBIC-L solution (10% w/w N-butyldiethanolamine, 10%w/w triton X-100, in H_2_O, Sigma Aldrich) and kept at 37 °C under continuous shaking. CUBIC-L solutions were refreshed every 5 days until the sample is cleared (~10 days). The cleared hemispheres were then transferred to CUBIC-RA solution (45% w/w antipyrine, 30%w/w N-methylnicotinamide, in H_2_O, Sigma Aldrich, refractive index=1.5) and again kept at 37°C under continuous shaking, with the CUBIC-RA solution being refreshed every 2 days until the hemispheres is completely transparent (~ 4 days).

### Microscopy

#### Confocal Imaging

To visualize fluorescent labeling of markers in brain sections, glass mounted brain sections were imaged with a Zeiss LSM700 upright confocal microscope with the ZEN Black image acquisition software (v2010, Zeiss), using the PLAN APO 20X objectives (0.8 NA, air) with the fixed wavelength 405nm, 488nm, 555nm, and 639nm lasers. Images were stitched with the acquisition software, further post-processed in FIJI (v2.0.0), and quantifications were conducted by experienced experimenters using the Adobe Illustrator (v2018).

#### Lightsheet Imaging

To visualize 3-D cleared hemispheres with monosynaptic inputs onto DMS ChAT neurons, CUBIC-L/RA processed anatomically labelled hemispheres were placed in a Zeiss Z1 lightsheet microscope with a PLAN APO 5X objective (0.16 NA, air) and the fixed wavelength 488nm and 561nm lasers to detect the fluorescently labeled mapping and autofluorescence respectively. The samples were imaged with a mounted PCO sCMOS camera in full frame (1920×1920 pixels) with an optical thickness of 12μm and exposure time of 200ms using the ZEN Black image acquisition software (v2010, Zeiss). The images were then first post-processed and fused at maximal intensity with Zen Lightsheet (Zeiss) and then stitched and visualized using the Arivis Vision 4D (v3.01, Arivis).

### Behavior

#### Fear conditioning

Mice were individually handled (5mins per day for 3 consecutive days) to habituate them to the experimenter. In the subsequent days following the last handling day, mice were trained (1 session/day) for 5 consecutive days. The animals were then placed into a plexiglass walled box (20cm wide, 22.5 cm long, 11.5cm tall) with an electrical gridded floor (Med Associates Inc) covered with black stripes (context A). Beddings were placed below. The bedding was replaced in between each animal and the box was cleaned between each animal with 70% ethanol. During each training session, the baseline contextual freezing activity was recorded for 2 minutes. Then 5 presentations of auditory conditioned stimulus (85dB, 7 or 12 kHz, counterbalanced) paired to an immediate 2s electric (0.6mA) shock (CS+) or without (CS−) each were presented for 10s in a randomized order, separated by a randomized inter-trial interval (ITI) with a duration between 50 and 150s. The animals were tested in the subsequent day following the last training day. They were placed into another plexiglass walled box (20cm wide, 25cm long, 15cm tall) covered with black polka-dots (context B) and cleaned with 1% acetic acid between each animal. The baseline contextual freezing to the new environment was also recorded for 2 minutes, following which, 4 instances of CS−and 4 of CS+ tones (85dB, 30s) were presented in sequential order separated with a randomized ITI with a duration between 70 and 130s. Videos were captured by a camera (The Imaging Source) from above at a fixed sampling rate (10Hz) and recorded mice were tracked with a behavioral acquisition and analysis software ANYMAZE (v5.23, Stoeling) that also controlled the delivery of tones (with Logitech speakers) and electric shocks (Med Associates Inc shock delivery box) with the TTL(time to live)-based master control box.

Freezing episodes lasting more than 1s were detected with the acquisition software and visually confirmed. The resulting freezing values and the timestamps of the CS presentations were exported.

#### Optogenetic manipulations

During the training days, the animals were restrained and the protruding optic fibers were bilaterally connected to custom-made optic cables (Thorlabs) that were connected via a commutator (Doric) to a 532-nm laser source (SUNON). The lasers are controlled by the behavioral acquisition software via a TTL master box that also controls the presentation of auditory tones and the delivery of electric shocks. The laser power was measured using the power meter (Thorlabs PM100A) and individually adjusted for each animal such that the power emitted from the optic fiber reaches 10mW. The lasers were turned ON continuously during the CS+ tone presentation throughout all five training days in both the control and experimental groups.

#### Calcium Imaging

On each day of behavioral experimentation, the miniscope mounted mouse was placed in the behavioral chamber and the midbrain calcium transients were recorded continuously with nVoke2 (Inscopix) at a 20Hz sampling rate. Simultaneously with the camera (The Imaging Source) that captures the behavioral movements and the acquisition software ANYMAZE (Stoeling) which also controls the presentation of auditory tones and the delivery of the electric shocks. Parameters such as the LED-power, the gain, and the electronic focus were adjusted on a mouse-to-mouse basis. The alignment of these recordings was made possible by a set of TTL triggers that were delivered by the behavioral acquisition software at the start and the end of the fear conditioning paradigm to the calcium imaging acquisition source. The behavioral paradigm included two pre-training sessions over two days where mice were presented to the contexts and tones in order to record a baseline activity to these cues. The mice were assessed for baseline contextual activity (context B in pre-training day 1 and context A in pre-training day 2) for 5 minutes and then to the two tones (30 seconds in pre-training day 1 and 10 seconds in pre-training day 2) three times each in a randomized order separated with a randomized ITI (between 50 and 150s in pre-training day 1 and between 70 and 130s in the pre-training day 2). In the day following the last pre-training day, the discriminative auditory cued fear conditioning paradigm was run as described previously with the exception that the CS− tone was pre-defined as the 7kHz tone and the CS+ tone pre-defined as the 12kHz tone. In addition to the resulting freezing values throughout the behavioral experimentation and the timestamps of the CS presentations, the recorded calcium transients (F/F) with respect to time were exported.

### Analysis

#### Freezing

The proportion of time mice spent freezing during the baseline contextual condition or during tone presentations were analyzed using custom-written Matlab (v2018b, Mathworks) scripts. Statistical comparisons in the proportion of freezing between the control and experimental groups and/or across experimental days were made and plotted using Prism (v8.2.1, GraphPad).

#### Calcium Imaging

Acquired calcium transients from the nVoke2 were first processed in the Inscopix Data Processing Software (IDPS, Inscopix), where the traces were pre-processed, spatially filtered, and motion corrected. Individuals cells were identified using the principle component (PCA) and independent component (ICA) analyses with no spatial or temporal down-sampling. Cells were visually verified by their maximal intensity projection visualizations and the placement of the region of interest (ROI) in these visualizations. Traces with abnormal physiological calcium transients (ie; transients lasting over minutes) were excluded. Time-stamped traces were exported to Matlab (v2018b, Mathworks), where custom-written scripts were used from there onwards. Calcium transients were first standardized (per trace per day), aligned to the presentation of external stimuli (tones, shocks) and then binned (500ms bins). Calcium activity surrounding the onset of tones (CS+ or CS−) were extracted (10s before and 45s after) and normalized to the baseline window (10s before tone onset). Average traces were generated and the average transients during the baseline period before the tone onset (10s) were compared to the initial period of tone presentation (also 10s). Cells were characterized either as excited (Wilcoxon rank-sum test p <0.01 and mean delta z-score > 0), inhibited (Wilcoxon rank-sum test p <0.01 and mean delta z-score < 0), or nonresponsive (Wilcoxon rank-sum test p >0.01). Shock responsive cells were characterized in the same manner by comparing the calcium activity 4s before CS+ presentation and 4s after the onset of the shock.

To examine whether cells could discriminate the presence of cues (CS−, CS+, or shock) or between cues (such as CS− from CS+), receiver operating characteristic analysis were used to calculate a discrimination index (DI, DI = (Area Under the ROC Curve – 0.5) × 2) from the distribution of activity during either cue presentation and during the corresponding baseline or from the distribution of activity during CS- presentation and the distribution of activity during CS+ presentation, respectively. The onset of cues was randomly shuffled across the trace (excluding the first 10s and last 14s) 1000 times and a distribution of randomized (DI)s were generated for each comparison. Cells with a DI that is outside of the 90% distribution of the randomized Dis were considered to be able to discriminate between the presentations of specific cue to baseline or to other cues.

To investigate whether the calcium activity pattern during the CS+ presentation is shaped towards the activity pattern of its paired aversive stimulus (shock), Euclidean distance (ED) as a measure of similarity is used and is calculated from the distribution of activity during first 4s of CS+ presentation and 4s following the shock onset for each pair across the training days. Again, the onset of the shock was randomly shuffled across the trace in a similar manner as previously mentioned and an average randomized ED is generated for each CS+/shock pairing per trace. Comparisons were made between the data-driven CS+/shock ED and randomized data-driven CS+/shock ED within and across training days.

#### Statistics

Statistical comparisons for the proportion of time spent freezing across days in the baseline contextual period or during tone presentations across different days were conducted with two-way analysis of variance (ANOVA) with multiple comparisons analyses following Bonferroni’s corrections (significance level = 0.05). Comparisons between behavioral discrimination to aversive paired and non-paired tones between control and experimental groups were done using unpaired t-tests (significance level = 0.05). In the analysis of the calcium imaging data, the comparison between calcium transients during tone presentations and time periods before were made with the Wilcoxon rank-sum test (significance level = 0.01).

## Acknowledgments

The authors thank the past and present lab members for constructive input and the team of the Imaging Platform of the Biozentrum. We are grateful to Wolf Heusermann for his technical support in brain clearing and light sheet microscopy. This work was supported by the Swiss National Science Foundation grant PP00P3_150683 and BSSGI0-155830.

## AUTHOR CONTRIBUTION

G.R. and K.R.T; Conceptualization. G.R, Z.L., N.H and K.R.T.; Methodology, Investigation; G.R, Z.L. and K.R.T Writing and Editing. K.R.T.; Funding, Supervision.

## Competing interest

The authors declare no competing interest.

## Figure Legends

**Figure S1: Midbrain neurons encode the association between CS+ and the foot-shock. (A)** Z-score heat-maps of average calcium transients. Black dashed lines depict the start (related to baseline) and end (related to shock) of the 4 sec time windows used for response classification. **(B)** Quantifications. **(C)** Z-score average curves of the calcium transients. **(D)** Z-score of the shock responses. **(E)** Discrimination index of the shock responsive cells. **(F)** Similarity index measured as Euclidean during C1. **(G)** Similarity index between the CS+ and shock responses during C3. **(H)** Eucleidean distance between CS+ and foot-shock responses during C5. **(I)** Similarity index between CS+ and shock responses across days and compared to randomized data points of the baseline calcium activity. **(J)** Legends. **(K)** Confocal images of midbrain sections showing GCamp6m-expressing neurons (green) and the lense track. The slices were counter-stained against TH (red).

**Figure S2: The sub-nucleus between the VTA and the SNc expresses pure neuronal cell types. A)** Confocal images showing mRNAs for tyrosine hydroxylase TH (red), Slc17a6 vesicular glutamate transporter-2 (cyan), Slc32a1 vesicular GABA transporter (green). **B)** Overall quantification. **C)** Quantification of TH+ cells.

**Figure S3: Pf thalamic cells provide monosynaptic excitatory inputs onto DMS-CINs. (A) (C)** and **(E)** Experimental designs. **(B)** Confocal image of the DMS (ChAT: magenta, Td-Tomato: red and synaptophysin: green). **(D)** Confocal image of DMS and the Pf. **(F)** Ih current and spontaneous firing of CINs. Light evoked excitatory current in CINs.

**Movie S1: Population activity in the midbrain during fear conditioning.** Example calcium transient observed under the field of view during the conditioning session 5.

**Movie S2: Input connectivity onto DMS-CINs.** 3D reconstruction of a ChAT-CRE mouse brain that was cleared after expression of rabies-EGFP in the DMS.

**Movie S3: Mouse behavior during optogenetic manipulation on conditioning day 1.**

**Movie S4: Mouse behavior during optogenetic manipulation on the test day.**

## References

1. Aosaki, T. et al. Responses of tonically active neurons in the primate’s striatum undergo systematic changes during behavioral sensorimotor conditioning. Journal of Neuroscience 14, 3969–3984 (1994).

2. Apicella, P. Tonically active neurons in the primate striatum and their role in the processing of information about motivationally relevant events. European Journal of Neuroscience 16, 2017–2026 (2002).

3. Zhang, Y.-F. & Cragg, S. J. Pauses in Striatal Cholinergic Interneurons: What is Revealed by Their Common Themes and Variations? Front Syst Neurosci 11, 80 (2017).

4. Brown, M. T. C. et al. Ventral tegmental area GABA projections pause accumbal cholinergic interneurons to enhance associative learning. Nature 492, 452–456 (2012).

5. Tan, K. R. et al. GABA neurons of the VTA drive conditioned place aversion. Neuron 73, 1173–1183 (2012).

6. Cohen, J. Y., Haesler, S., Vong, L., Lowell, B. B. & Uchida, N. Neuron-type-specific signals for reward and punishment in the ventral tegmental area. Nature 482, 85–88 (2012).

7. Root, D. H., Estrin, D. J. & Morales, M. Aversion or Salience Signaling by Ventral Tegmental Area Glutamate Neurons. iScience 2, 51–62 (2018).

8. de Jong, J. W. et al. A Neural Circuit Mechanism for Encoding Aversive Stimuli in the Mesolimbic Dopamine System. Neuron 101, 133–151.e7 (2019).

9. Morales, M. & Margolis, E. B. Ventral tegmental area: cellular heterogeneity, connectivity and behaviour. Nat. Rev. Neurosci. 18, 73–85 (2017).

10. Klug, J. R. et al. Differential inputs to striatal cholinergic and parvalbumin interneurons imply functional distinctions. Elife 7, (2018).

11. Matsumoto, N., Minamimoto, T., Graybiel, A. M. & Kimura, M. Neurons in the thalamic CM-Pf complex supply striatal neurons with information about behaviorally significant sensory events. Journal of Neurophysiology 85, 960–976 (2001).

