## Supplementary figures and images for "Midbrain modulation of conditioned learning"

### Supplementary Figure 1

#### Rizzi*, Li* et al. Figure S1

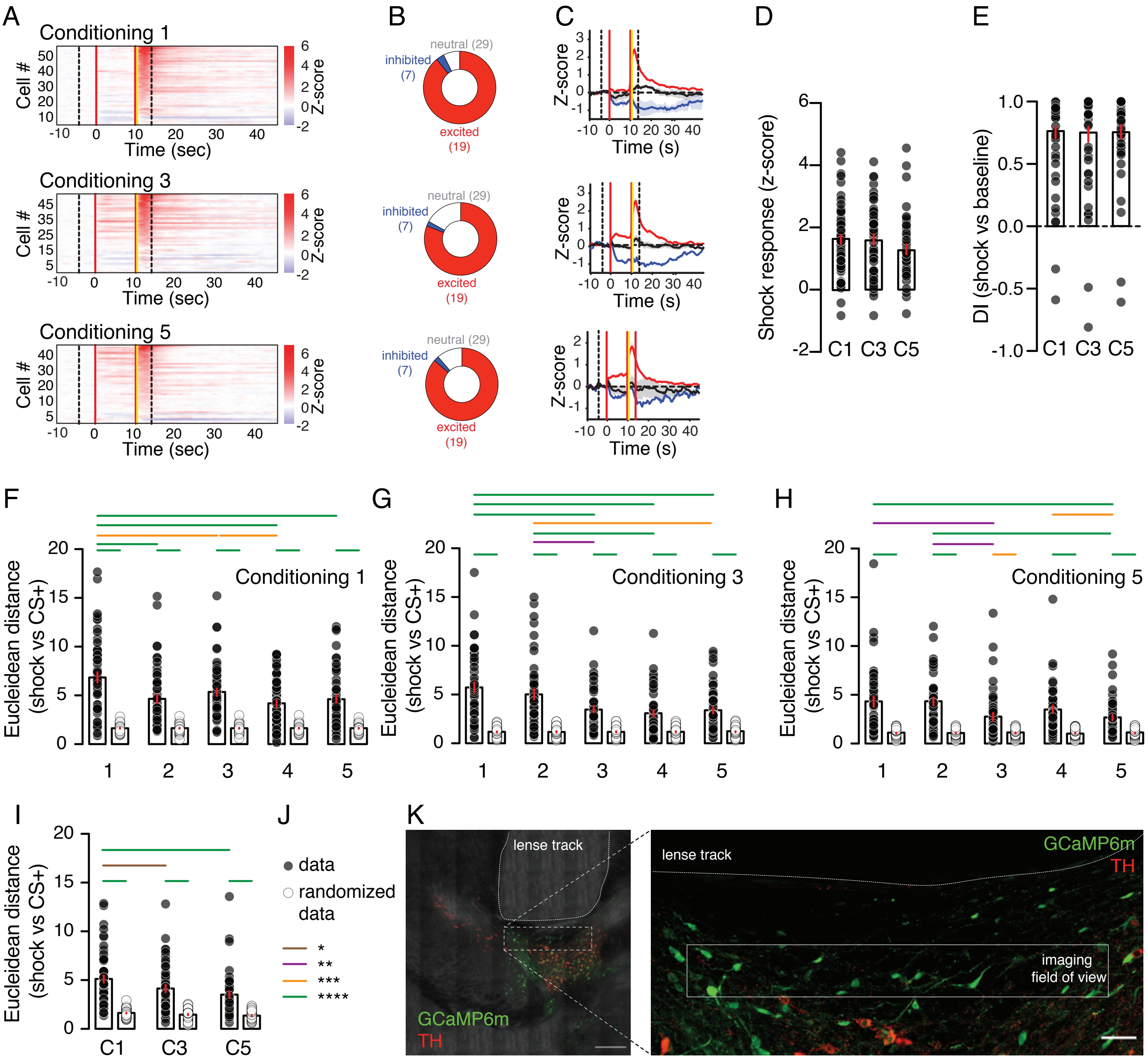

### Supplementary Figure 2

Rizzi*, Li* et al. Figure S2

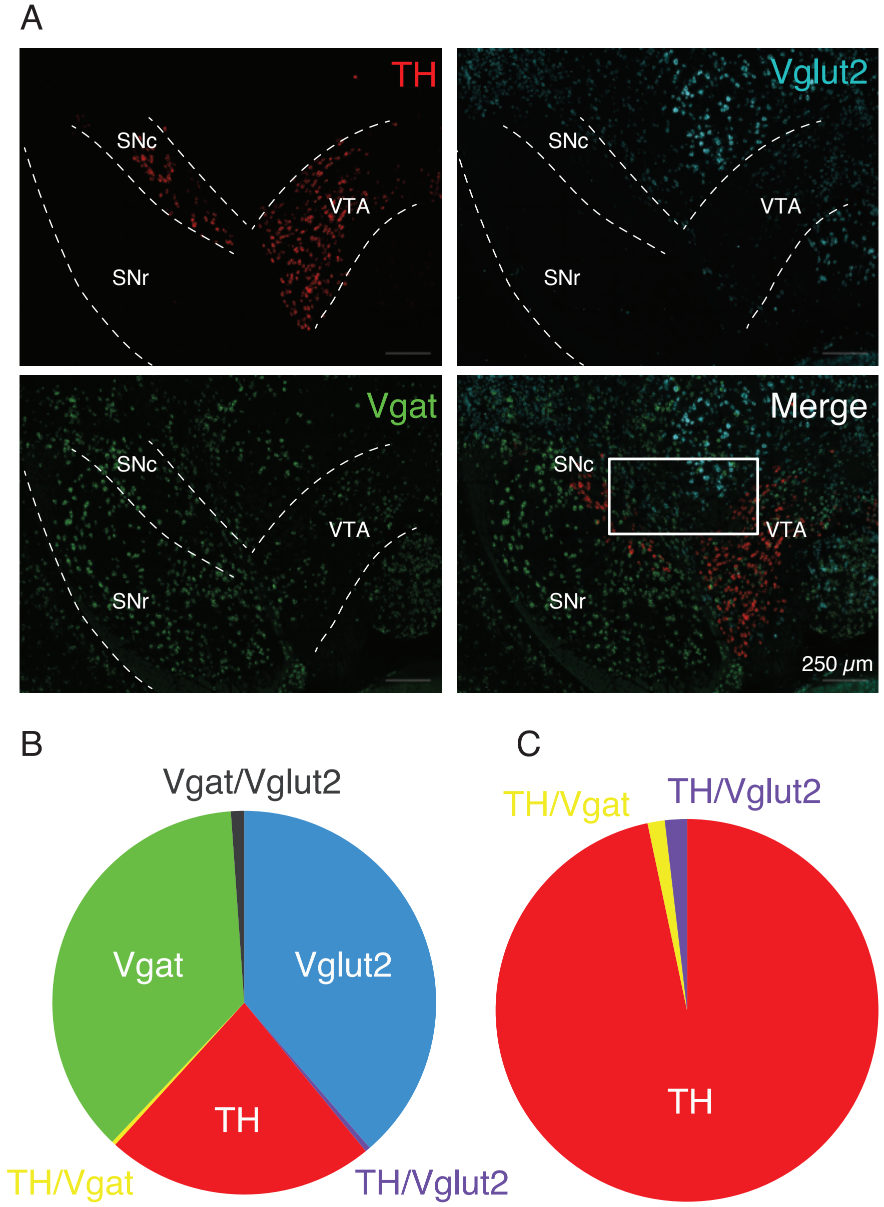

### Supplementary Figure 3

Rizzi*, Li* et al. Figure S3

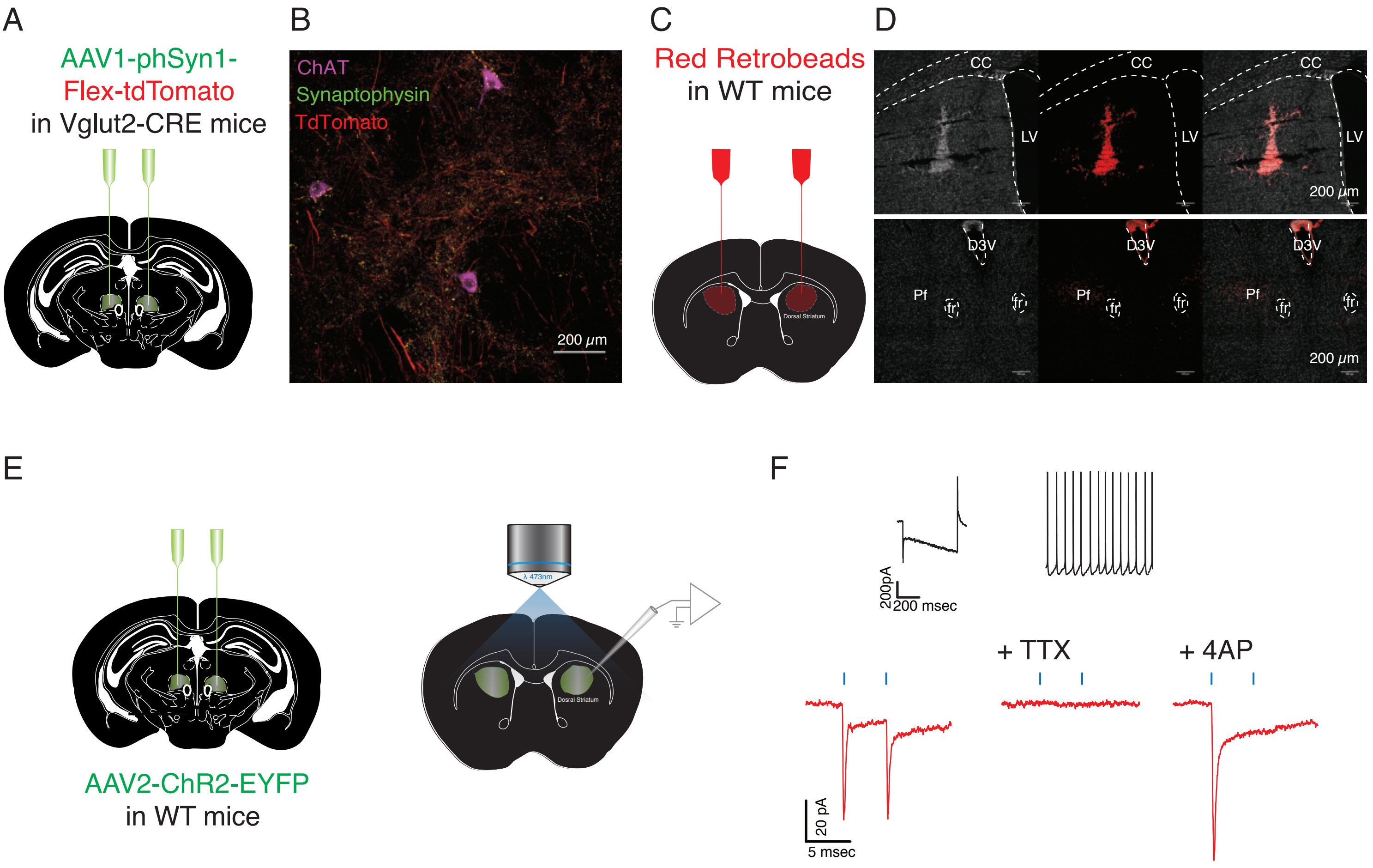
